# HtPIP: High-Throughput Phage Isolation Platform increases diversity and reduces isolation time using multiple bacteria

**DOI:** 10.64898/2026.03.10.710857

**Authors:** Ben Diaz, Tessa House, Meghana Padala, Joseph S. Schoeniger, Catherine M. Mageeney

## Abstract

Bacteriophages are ubiquitous in nature, but relatively few have been isolated and characterized compared to the number of bacterial strains. Phage biotechnology applications benefit from a diverse library of isolated phages to kill or transfer genetic material to a bacterium of interest. However, scaling phage discovery for diverse bacterial hosts can be time consuming and costly. We developed an approach to capture novel phages for multiple bacteria strains in parallel from an environmental sample using commercially available 0.2-micron filter plates. Using this **H**igh-**t**hroughput **P**hage Isolation **P**latform (HtPIP), we isolated twelve novel phages spanning nine diverse bacterial host genera. Eleven of the isolated phages define new phage species with nine also defining new genera. We show the HtPIP can discover both DNA and RNA phages; including a *Tectiviridae* infecting *Pseudomonas putida* mt-2 and a *Leviviricetes* infecting a *Microbacterium* isolate, which represents the first cultured RNA phage infecting a host outside of proteobacteria. Using a metagenomic approach, we demonstrate that the HtPIP captures a higher proportion of novel phages compared to traditional low-throughput methods.

## Introduction

Bacteriophages (phages) are viruses that infect bacteria. They are found in almost every environmental niche on Earth that supports bacterial life, including wastewater (1), soil (2), the ocean (3), and mammalian microbiomes (4). Most phage have access to two lifecycles which allow the bacteria/phage relationship to be predatory when they complete the lytic lifecycle, or mutualistic during the lysogenic lifecycle (5-7). Additionally, phages can carry novel auxiliary metabolic genes to enhance host metabolism (8), or genes that enhance host pathogenesis or predator resistance. The diversity of phages in the biosphere allow for a wide range of applications in biotechnology as gene delivery vehicles (9), sources of new synthetic and molecular components (10, 11), as biocontrol agents to prevent plant disease (12, 13), agents to combat biofilms (14), and as therapeutics for antibiotic resistant infections in humans and animals (15, 16).

Most bacterial strains do not have isolated phages known to infect them available for biotechnology applications involving phages. Many applications typically require a cocktail comprised of several complementary phages (15) to prevent prompt phage resistance, further aggravating this gap in availability. It follows that for many phage-based applications to be successful, rapid discovery methods or a phage bank with numerous diverse phages for each bacterial host is required. Traditional phage isolation is, however, low throughput and costly for multiple hosts, and thus does not always yield the quantity nor diversity of phages needed for a phage bank (17, 18).

A typical process for isolating phages from the environment is to add the environmental sample (soil, water, sewage) to a culture of a single bacterial host of interest, incubate, and then recover phage particles and remove bacteria (typically through filtration). The filtrate is then tested for presence of phages for the bacteria of interest (18) typically by plaque assay. While methods for higher-throughput phage discovery have been developed in recent years by scaling this process for a 96 well plate (19) they can remain time-consuming and costly due to multiple manual filtration steps. Inspired by the iChip, which enables culturing of previously uncultivable bacteria by growing them within a compartment enclosed by semipermeable membranes within an environmental sample (20), we sought to design a high-throughput method that captures a higher diversity of native phages by allowing low abundance phages to bloom alongside the native bacteria living in the environmental samples (21). In addition, we aimed to reduce the labor-intensive and potentially hazardous steps of centrifugation and filtration of environmental samples (22, 23).

We developed and tested the feasibility of a **H**igh-**t**hroughput **P**hage **I**solation **P**latform (HtPIP). We sought to establish a robust method with commercially available materials to ensure reproducibility and reduce phage isolation time for multiple bacterial strains concurrently. Using HtPIP, we isolated twelve novel phages infecting nine bacterial hosts. These include diverse morphotypes (siphoviridae, myoviridae, tectiviridae, and leviviridae) and many new genera and species of phages. Furthermore, we isolated the first *Leviviricetes* infecting a gram-positive host (*Microbacterium*), demonstrating the utility of HtPIP to capture both DNA and RNA viruses. We then explored the potential diversity of the metavirome captured from wastewater influent incubated with *E. coli* and found that a higher proportion of novel and diverse phages are captured with this HtPIP versus traditional low throughput methods.

## Materials and Methods

### 2.1 Bacterial Cultures

Bacteria were grown overnight shaking at 200 RPM at 20°C. 20°C was chosen to allow all bacteria to grow and be infected by phages, as *Pseudomonas putida* has been shown to defend against phage infection at 37°C (24). The following bacteria were grown in LB media (Sigma Aldrich): *Escherichia coli* MG1655, *Pseudomonas putida* mt-2 (ATCC 33015), *P. putida* S12, *Bacillus subtilis* NCIB 3610, and *Burkholderia cenocepacia* K56-2 (obtained from BEI NR-20535). The following bacteria were grown in R2A media (Teknova): *Variovorax* sp. SCN, *Variovorax* sp. OAS795, *Rhodococcus rhodochrous* 372 (ATCC 13808), *Rhodococcus qingshengii* S10, *Rhodococcus* sp. MSC1_016, and *Sphingopyxis fribergensis* MSC1_008. *Microbacterium* sp. was grown in Difco ISP media (BD). All cultures were supplemented with 1 mM CalCl_2_ (Sigma Aldrich).

### 2.2 Environmental Sample Collection

Samples were collected from the sites listed in **Table 1**. Untreated wastewater influent samples were collected with permission by Livermore Water Reclamation plant in Livermore, CA, USA. 250 mL of liquid samples were collected from wastewater and pond sources, while approximately 100 mL volume of solid samples were collected from each solid source.

**Table 1.** Sampling sites used in this study. Samples were collected from a diverse set of sites in the San Francisco East Bay, USA and agricultural field site in Washington State, USA. The sample name, GPS coordinates, and state of the sample (liquid/solid) are listed.

| Sample | GPS Coordinates | Sample State |
| --- | --- | --- |
| **Wastewater Influent 1, 2, 3** | 37.69073324466393, -121.80667889554306 | Liquid |
| **Tomato rhizosphere** | 37.68343382977756, -121.74488605243637 | Solid |
| **Compost - backyard** | 37.93132569198107, -121.7123907950711 | Solid |
| **Topsoil** | Purchased from home improvement retailer | Solid |
| **Compost - community center** | 37.677616812367795, -121.73652562520904 | Solid |
| **Washington State University field site in Prosser, WA** | 46.252480744413305, -119.73798155092547 | Solid |

### 2.3 Traditional phage isolation

Traditional enrichment filtrates were processed and filtered the same day HtPIP was started. Liquid environmental samples were filtered through a 0.2-micron filter (Millipore Express PES Membrane). SM buffer without gelatin (150 mM NaCl (Sigma Aldrich), 40 mM Tris-Cl (pH 7.4) (Teknova), and 10 mM MgSO4 (Sigma Aldrich)) was added to solid samples to make a slurry, inverted gently several times, centrifuged for 30 minutes at 5000 RCF, and the supernatant was filtered through a 0.2-micron filter. Enrichment cultures were set up with 1 mL of filtrate, 4 mL of fresh LB liquid supplemented with 1 mM CaCl_2_, and 50-1000 μL of an overnight bacterial culture. Overnight cultures of faster growing bacteria, (*E. coli* and *P. putida, B. subtilis)*, were added at lower ratios (50 μL: 1:80 dilution), while all other bacteria used were added at higher ratios (1000 μL: 1:4 dilution). These mixtures were grown for ∼68 hours at 20°C shaking at 200 RPM, then filtered again through a 0.2-micron filter. We used double agar overlays to determine if phages were present for hosts of interest by mixing 250 μL enrichment filtrate, 50-1000 μL of overnight bacterial culture and 3 mL’s of LB top agar (0.5% agar, 1 mM CaCl_2_) and plated onto LB Agar plates supplemented with 1 mM CaCl_2_.

### 2.4 HtPIP phage isolation

Freshly diluted bacterial cultures at the same ratio described above (1:80-1:4) were loaded into a MultiScreen-GV Filter Plate (Millipore Sigma, MAGVS2210), with the underdrain removed, before placing on top of a soil slurry made with SM buffer or unfiltered wastewater. HtPIP plates were incubated 20°C, at 100 rpm for 68 hours. A separate plate was used to filter the subsequent enrichment cultures with the underdrain intact. Liquid samples were processed by transferring 150 μL from each well to a new filter plate placed on top of a 96 well plate (Nunc 96-Well Polystyrene Round Bottom Microwell Plates – 268200) and centrifuged for 10 minutes at 2000 RCF. Semi-solid samples (0.5% agar with growth media (i.e. LB, ISP, R2A), melted and kept molten at 55°C) wells were processed by adding 100 μL of phage buffer to each well, pipetting up and down several times slowly, then transferred to the clean filter plate. If the sample did not filter, centrifugation was repeated up to 2 more times.

### 2.5 Phage Purification of Isolated Phages

In samples where plaques or clearing of lawns were observed, 2 individual plaques were picked, resuspended in 100 μL SM buffer, and diluted to 10^-5^. These dilutions were mixed with fresh overnight bacterial culture diluted in each host’s respective molten top agar media (LB, R2A or ISP). This process was repeated for three purifications total, wherein lawns were flooded with 8 mL SM buffer, scraped, centrifuged briefly (500 RCF for 5 minutes) and the supernatant was filtered through a 0.2µM filter. Genomic DNA was extracted from these purified lysates using Norgen Phage DNA Isolation kits without modification. Libraries were prepped with the Illumina DNA Prep Tagmentation kit and Illumina Unique Dual Indexes.

For phages where the Norgen kit did not yield high quality DNA, lysates were processed by SeqCoast Genomics (Portsmouth, NH, USA) where samples were lysed using MagMAX Microbiome bead beating tubes, and DNA was extracted using the Qiagen DNeasy 96 PowerSoil Pro QIAcube HT Kit (Listed **Supplementary Table 1**).

*Microbacterium* phage Later RNA was extracted using the Monarch Mag Viral RNA/DNA extraction kit (New England Biosciences). Libraries were prepped using the Illumina Stranded Total RNA Prep with Ribo-Zero Plus kit without modifications and sequenced on an Illumina NextSeq 2000.

All DNA libraries were sequenced using Illumina sequencing platforms (see **Supplemental Table 1** for platform).

### 2.6 Phage Metavirome Collection

Metaviromes were generated from traditional and HtPIP methods. The same overnight culture of *E. coli* MG1655 was diluted to the same concentration in liquid (50 μL overnight culture/mL sterile LB supplemented with 1 mM CaCl_2_) and loaded onto a filter plate “HtPIP Liquid” or added to 1 mL LB top agar (“HtPIP Semi-Solid”), or aliquoted into a tube with 1 mL 0.2µM filtered wastewater (“Traditional”). All HtPIP samples were sealed with Microseal “B” PCR Plate sealing film (Bio-Rad) and placed on top of 100mL unfiltered wastewater. All samples were incubated at 20°C, at 100 rpm for 68 hours.

Enrichment cultures were filtered using a 0.2µM syringe filter (“Traditional”) or via centrifuge (“HtPIP”), and then serially diluted and combined with freshly diluted LB top agar and plated using the double agar overlay technique on LB agar plates. After overnight incubation at 20°C, phage buffer was added to the plates, and the top agar was gently scraped. The top agar/phage buffer slurry was centrifuged for 10 minutes at 4000 RCF, and then 0.2-filtered using a syringe filter.

DNA was extracted from 1 mL of each enrichment lysate using a phenol-chloroform extraction. Before extraction, Proteinase K (New England Biolabs) and 0.05%SDS (Sigma Aldrich) were added, and the lysates were incubated at 55°C with gentle shaking every 15 minutes. Next, an equal volume of phenol – chloroform – isoamyl alcohol mixture (25:24:1) was added to the samples and incubated at room temperature for 10 minutes. Samples were spun at 4°C at 10000 RCF for 10 minutes, and the aqueous phase was removed from the top. An equal volume of chloroform was added to the aqueous layer, incubated at room temperature for 10 minutes, centrifuged at 4°C at 10000 RCF for 5 minutes, and the aqueous phage removed to a new tube. The chloroform extraction was repeated twice. The resulting aqueous extractions were then precipitated with PEG (final concentration 8X%) and NaCl (final concentration 1 M). This was let sit at room temperature for 30 minutes and then centrifuged at 10000 RCF at room temperature for 30 minutes. The pellet was washed once in 100% and twice in 70% ice-cold ethanol before drying for 15 minutes at 37°C. The pellet was resuspended in deionized water and measured with QuBit before library preparation. Libraries were prepared with the Nextera XT kit and sequenced for short-read Illumina sequencing at Genewiz (South Plainfield, NJ).

### 2.7 Isolate phage assembly and annotation

Raw sequencing reads for both metavirome experiments and isolated phages were processed with BBDuk (https://sourceforge.net/projects/bbmap/) with the following parameters specified (ktrim=r, k=21, mink=11, hdist=1).

For isolate phage sequences, other than *Microbacterium* phage Later, trimmed high quality reads were assembled using spades (v3.9.0)(25) with the standard parameters for paired-end sequencing. The resultant spades assembly was further filtered to remove small (<1000bp) and low coverage assemblies (<1% max coverage). Final phage genomes were annotated using Prokka (v1.11) (26) for open reading frame (ORF) calling and our DiMER pipeline (https://github.com/sandialabs/DiMER) for functional prediction. Resultant phage genome annotations were manually checked for any large gaps and to resolve translational frameshifts. tRNAscan-SE (v2.0.12) (27) was used to predict t(m)RNA genes in our phage genomes using the following additional parameters: -G -I. Phage genome statistics are listed in **Supplementary Table 1**. Phage taxonomy was predicted using the web-based version of taxMyPhage (28) and vContact2 (v0.9.19) (29) at KBase (30). Phage genome maps were created with web-based Proksee (31). CheckV (32) (end_to_end, -t 5) was used to determine completeness. The nucleotide sequence for each phage was input as the query using the core nucleotide database (core_nt) using web-based BLASTN to determine related phages. We used the web-based VIPTree (33) and VIRDIC (34) to explore any evolutionary relationships with published phages.

Two clonal isolates of *Microbacterium* phage Later were sequenced. Raw reads from each were processed with BBDuk as described above. The trimmed high-quality reads were assembled using MEGAHIT (v1.2.9) (35, 36) with standard parameters for paired-end sequencing. The resultant assemblies yielded multiple contigs (clone 1: 38 contigs and clone 2: 68 contigs) with few larger than 1000bp (3 contigs and 9 contigs, respectively). We used BLASTN to determine if these contigs were bacterial or viral. Only one contig in each assembly aligned with a viral sequence (Riboviria sp. [OQ424403.1]). These two contigs from the *Microbacterium* phage Later clonal isolates were aligned to each other and showed 99.9% identity over 100% of the contig. The resultant contig was annotated as described above with Prokka. We sought to understand if predicted orf3 was the lysis protein in *Microbacterium* phage Later. The protein sequence for the ORF3 was input into the web version of DeepTMHMM (37).

### 2.8 Metavirome assembly and clustering

Trimmed high quality reads from our metaviromic samples were assembled using MEGAHIT (v1.2.9) (35, 36) with standard parameters. We used the MVP pipeline (38) modules 00, 01, 02, 03, 04, and 05 to cluster resultant viral contigs and obtain protein sequences for each viral contig as individual samples and across the entire dataset. VContact2 (39) (version 0.11.3) with default settings was used to cluster viral contigs. Ggraph was used to visualize clusters.

### 2.9 Transmission electron microscopy

We sent fresh high titer lysates for TEM imaging on a Hitachi HT7800 by Tagide deCarvalho, Keith R. Porter Imaging Facility, University of Maryland, Baltimore County.

## Results

### 3.1 Development and Optimization of HtPIP

Phage discovery using a semipermeable membrane plate arrayed with multiple bacterial strains was motivated by two goals: to capture a wider diversity of phages for a diverse set of hosts and to spend less time processing environmental samples. We were inspired by the iChip system, which allows for cultivation of diverse bacteria from the environment by allowing bacteria to exchange nutrients with their native habitat through a semipermeable membrane (20). We reasoned in environmental samples diverse phages can bloom alongside bacteria *in situ*, and we would capture a wider variety of phages by tapping into this dynamic *in situ* pool. The parallel screening is aimed to increase the rate of success of finding a phage compared to traditional methods in two ways: 1) Given the inherent hit-or-miss during phage discovery for a specific host, screening diverse hosts in parallel increases the odds of capturing phages for some; 2) solid samples can be inhomogeneous, and phage may bloom at very low levels, so spatially distributed replicate wells containing the same host may increase the odds of capturing diverse phages for that host. Additionally, we aimed to remove the step of centrifuging and filtering environmental samples, which can be difficult and time consuming especially for soil samples (22).

To ensure that this experimental system (**Figure 1**) was sufficient to meet these goals, we tested several fundamental properties critical for phage discovery, including phage transport through the 0.2-micron membrane, bacteria seeding density, media, and plate sealing. We utilized a model host-phage system (*E. coli* MG1655 and *coliphage* Lambda) to optimize HtPIP. We tested an array of conditions to determine which resulted in the highest yield of phage after enrichment: liquid vs semi-solid (0.5% agar), host density: low (10^5^ CFU/mL) vs high (10^7^ CFU/mL), sealing: ambient airflow (hard plastic cover) vs low airflow (Microseal adhesive B cover) (**Figure 2a**). Lambda was diluted in phage buffer to 1×10^6^ PFU/mL inside the sterilized sample reservoir and the filter plate was loaded with *E. coli* MG1655 and multiple negative controls (phage buffer, LB broth, top agar). The loaded plate was placed on top of the Lambda suspension, shaken for 1 hour at 50 RPM at 20°C (room temperature), and let incubate overnight at 37°C. The next day liquid wells were pooled and filtered by hand via syringe 0.2 µM filter, while top agar cultures were mixed with 200 μL of phage buffer, pooled, and then filtered by hand via syringe 0.2µM filter.

**Figure 1.**
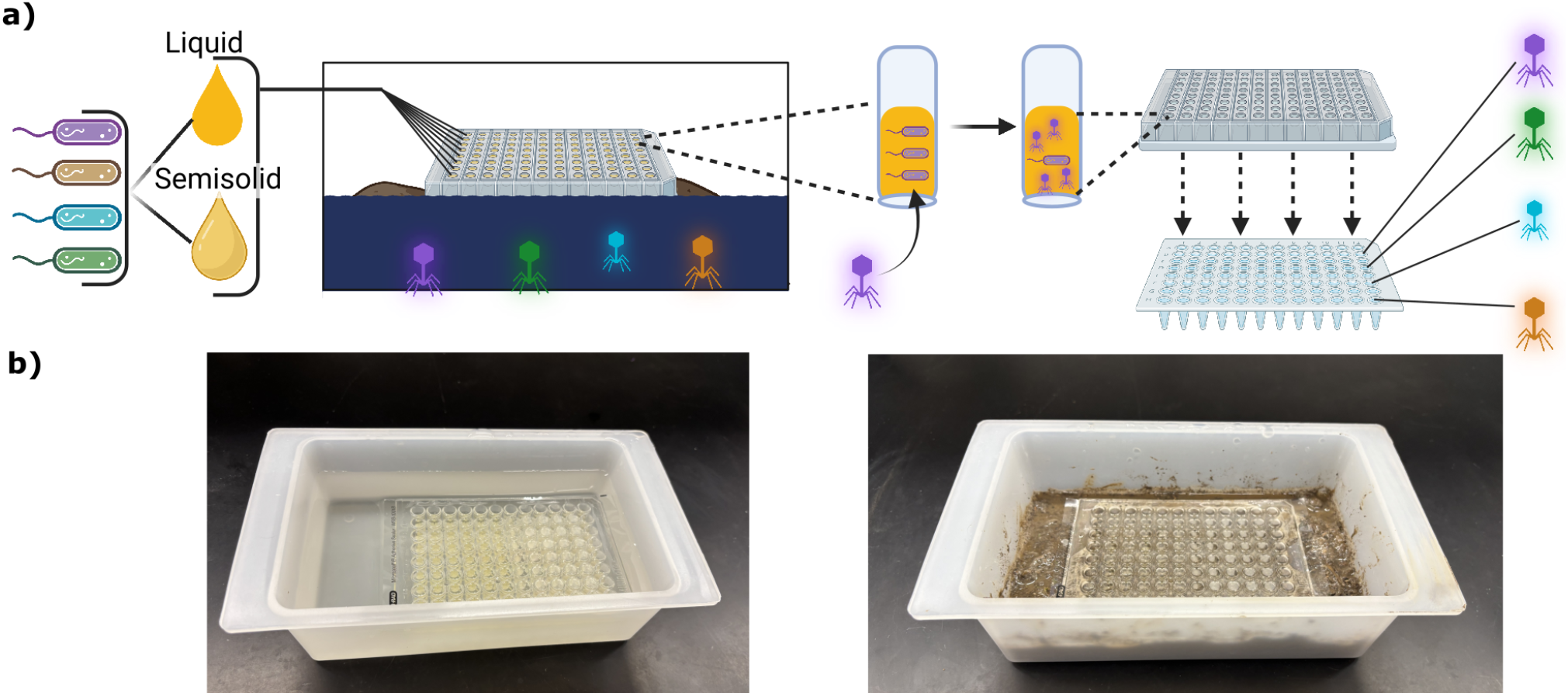
HtPIP setup and workflow. **a)** Schematic of process. Bacteria are diluted into fresh liquid or semisolid media, then aliquoted into a semipermeable plate. The plate is placed on top of a soil slurry or liquid sample, where phages can percolate through the membrane, but bacteria are contained. After incubation, the wells are filtered, and phages are recovered from the filtrate. **b)** Photo of plate incubating on top of a liquid (left) or soil slurry (right).

**Figure 2.**
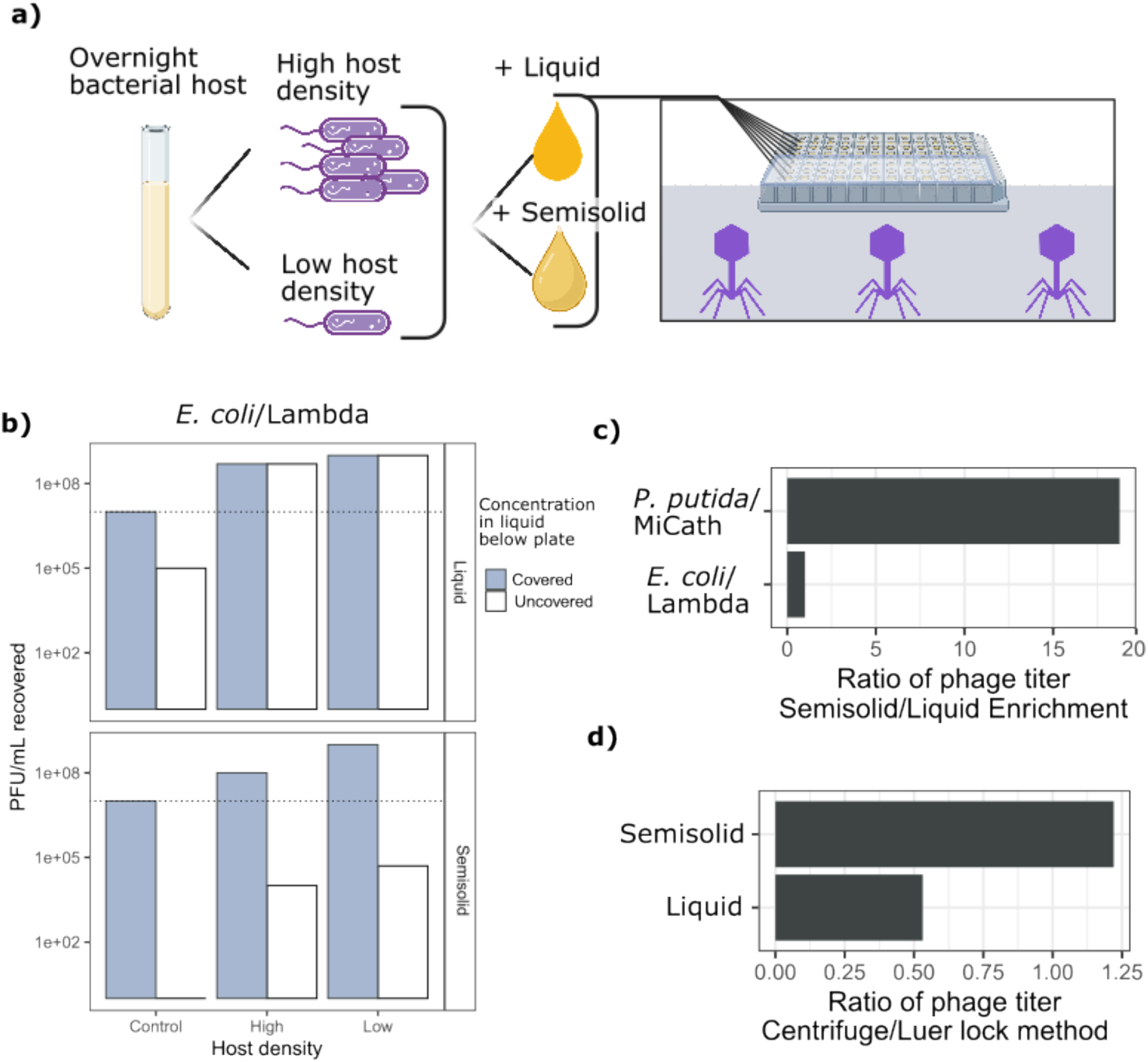
Optimization of phage isolation conditions using HtPIP. **a)** Schematic of HtPIP conditions used for optimization with known phage concentrations in the reservoir below the plate. **b)** Experiments with *E. coli/lambda* combination, testing the effect of covering half of the plate with a PCR plate sealer (“Covered”/”Uncovered”), starting host density for enrichment (“High”/”Low” + control – only sterile media added to wells), and different enrichment media (“Liquid” –LB broth/ “Semisolid”-0.5% LB Agar). The dashed line represents the concentration of phages that were present in the liquid below the plate – a 1:1 ratio in the control means effectively all of the Lambda phage particles passed through the filter. **c)** Testing the effect of enrichment media with a non-model host/virus combination (*P. putida*/MiCath) on phage titer captured with the HtPIP (covered plates). Low starting host densities were used for both host/virus combinations. **d)** Testing the effect of filtering method after enrichment with *P. putida*/MiCath with the HtPIP (covered plates). Centrifuge filtering saves time by filtering the entire plate at once, while syringe filtering is a typical method used to filter each enrichment lysate by hand.

In our lambda experiments we found that phages could percolate from the sample reservoir through the membrane to infect bacterial hosts, since the negative control had ∼100% recovery of the phages (**Figure 2b**). We learned that a sealed top was necessary to capture phages when using semisolid media and significantly increased titers in liquid conditions (**Figure 2b)**. It appeared that wells with lids left loose had less overall liquid, even in the media only control wells after incubating overnight. This may be a sign that the sealed plates prevent evaporation, or they prevent liquid equilibrating with the outer liquid level, which is slightly lower than the liquid cultures growing inside the plate. Lastly, we saw a modest increase in phage titer with low starting host density in semisolid plates and no change dependent on host density in phage recovered from liquid conditions (**Figure 2b**).

We next wanted to ensure the utility of our HtPIP plate for non-model bacteria and phages. We used as a non-model phage-host system *P. putida* S12 and phage MiCath (40). MiCath was 18X more concentrated when harvested from a semisolid enrichment compared to liquid enrichment **(Figure 2c)**. Coliphage lambda did not have the same preference for semisolid enrichment, with an equal ratio of phages recovered from liquid and solid covered plates (1X, **Figure 2c**). We also sought to test the effect of centrifuge filtering versus manual syringe filtering to ensure complete capture of phage particles, removal of bacteria, and no sample type bias existed. Both liquid and solid enrichment cultures were able to be filtered and yielded a recovery within a 2-fold range (50% and 125% of hand filtering), of the concentration of phages compared to manual filtering with a syringe and luer-lock filter (**Figure 2d)**. We considered a two-fold difference to be within acceptable range of hand-filtering since it saves time and limits cross contamination. These experiments gave us optimized conditions to search for phages in a variety of environmental samples. We recommend a sealed plate top, inoculation with low host densities, and filtration by centrifugation for the highest yield of phages from environmental samples.

### 3.2 HtPIP enhances discovery of novel phage for multiple bacterial hosts

Using our optimized HtPIP (covered plate, low host density, centrifuge filtering, both liquid and semi-solid), we tested a diverse set of environmental samples (**Table 1**) to discover new phages for a large phylogenetic range of bacterial hosts. Since semi-solid and liquid enrichments may capture different phages, so both were used in future experiments. Overall, we tested 11 bacterial strains against 9 different environmental samples (**Supplementary Table 2**). Phage plaques were observed in 31% (15/48) of the phage/host/sample combinations. We further purified and sequenced a subset of these phages (**Supplementary Table 1**). Using the HtPIP, we discovered 12 novel phages across 9 phylogenetically diverse bacterial host strains. We also recovered *Pseudomonas putida* S12 phage MiCath from the same compost sample it was originally isolated from (40). The new phages can be categorized into three taxonomic groupings: 1) maps to known species (Tolp), 2) maps to known genus (EW1 and Eliverwaste), 3) defines new genus and species (PLiverWaste, Gard, SweetSinh, Perlinasted, Prosser1, Prosser2, Maizie, Soil63, Later). The large number of phages that represent new phage genera highlights that the HtPIP can recover novel, highly diverse phage for non-model hosts.

Regarding the three phages that have close relatives, none of these phages are identical to any previously isolated phage (genome statics in **Supplementary Table 1**). Tolp is very similar to other *tectiviridae* (97.74% ID over 100% genome to PRD1 [NC_001421.2]) and is a member of the *Alphatectivirus* PRD1 species. EW1 and Eliverwaste define novel species within their respective genera. EW1 has similarity to T1 and is most similar to vB_Henu5_2 [PQ362312.1] (90.56% ID over 95% genome). EW1 also carries the cor superinfection exclusion protein which is used to prevent subsequent phage infection (41) (**Figure 3A**). Eliverwaste belongs to the *Felixounavirus* genus and is most similar to JLBYU32 [OK272490.1] (98.54% ID over 93% genome). Eliverwaste contains multiple tRNA genes and a ribonucleotide pathway (**Figure 3A**).

**Figure 3.**
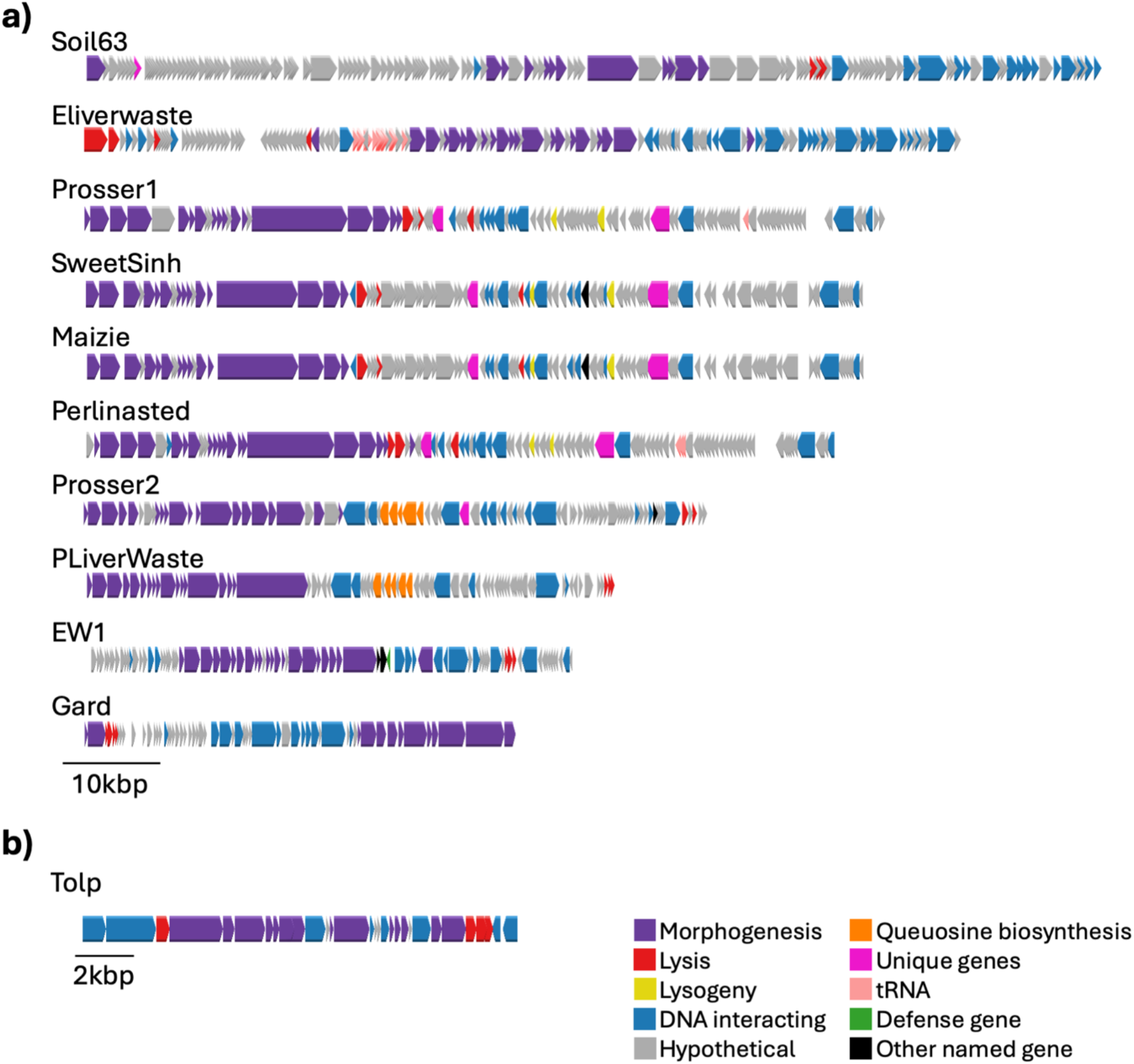
Genome Maps of Isolate Phages discovered using HtPIP. Each phage genome is shown, and genes are colored according to function. **a)** shows all phages larger than 40kbp ordered by size. **b)** shows phages smaller than 40kbp. A bar indicating size is below each series of maps.

Of the remaining nine phages that were novel and define new genera and species of phages, four are new phages for three *Rhodococcus* strains (**Supplementary Table 1**). Perlinasted was isolated on *R. qingshengii* S10 and resembles a previously isolated phage from our group Perlina (PP782351.1; 97.43% identity over 98% genome) (42). Perlinasted contains 3tRNA genes (**Figure 3A**). SweetSinh and Maizie were both isolated on *R. rhodochrous* 372. They are 99.8% identical to each other, suggesting they are the same phage species, and have similarity to *Rhodococcus* phage Reynauld (OR159659.1; 80.3% over 10% of genome). Prosser1 was isolated on *Rhodococcus* sp. MSC1_016, an isolate from a novel soil consortium MSC-2 (43). Prosser1 shares little similarity with any phages at NCBI with the closest relative is *Rhodococcus* phage Mbo2 (ON191531.1; 78.75% identity over 1% of genome) and encodes a single tRNA gene. All *Rhodococcus* phages discovered contain a cas4 exonuclease and CobT-like cobalamin biosynthesis protein. All four phages are temperate phages encoding a tyrosine integrase (**Figure 3A**).

Two new *Variovorax* phages were discovered, Gard, isolated on *Variovorax sp*. SCN, and Soil63, isolated on *Variovorax sp*. OAS795. Gard is similar to *Variovorax* phage VAC51 (OX359471.1; 81.61% ID over 69% genome). Soil63 is a completely novel phage with only 228bp homology found in another phage at NCBI (Rhizobium phage RHpl_I4 [MN988552.1]). Soil63 is the largest phage (103.3kbp) discovered thus far using our HtPIP. Soil63 encodes a dCTP deaminase, which is typically classified as an auxiliary metabolic gene (44). Overall, due to the lack of sequence conservation, only 22.7% of genes were able to be assigned a function (32/141) (**Figure 3A**).

Although there have been many recent phages discovered for *P. putida* (24, 40), PLiverWaste is novel only sharing 73.81%ID over 2% of the genome with *Pseudomonas* phage Baskent P4_1 (PP992516.1). PLiverWaste contains a queuosine biosynthesis pathway, suggesting the PLiverWaste DNA is protected from restriction endonucleases (40, 45).

We discovered a new *Sphingopyxis* phage Prosser2. This has similarity to a previously isolated *Sphinngomonas* phage vB StuS MMDA13 (NC_072503.1; 78.22%ID over 66% genome). Prosser2 has typical genes expected in a phage genome and a queuosine biosynthesis cassettes which has been shown to modify phage DNA to prevent restriction by the host (40, 45).

To further characterize each of the eleven DNA phages above, we used TEM to determine the phage morphotype (summarized in **Supplementary Table 1**). Prosser2, Soil63, PLiverWaste all have typical siphoviridae morphology (**Figure 4a**). These phages all have a 1:3-4 head to tail ratio. The *Rhodococcus* phages (SweetSinh, Maizie, Perlinasted, and Prosser1) all have similar siphoviridae morphology with very long tails (400-460nm) resulting in a head to tail ratio of 1:7 (**Figure 4b**). We discovered two myoviridae, Gard and ELiverwaste (**Figure 4c**). Lastly, we confirmed the *tectiviridae* morphology for Tolp (**Figure 4d**).

**Figure 4.**
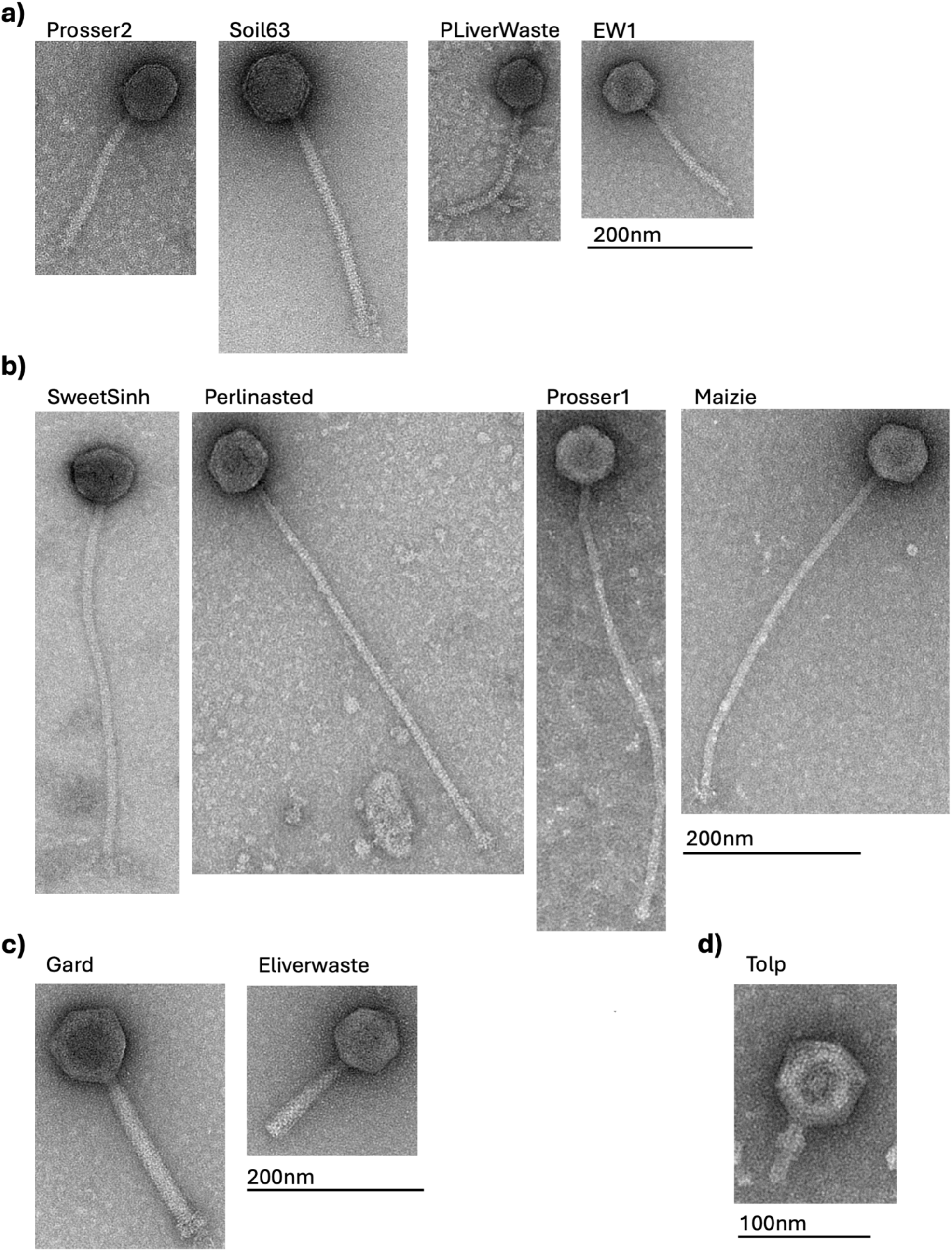
TEM images of all phages found using HtPIP. Here we show TEM images. We have grouped the phages by morphotype. **a)** Phage with siphoviridae morphology, **b)** long tailed siphoviridae, **c)** myoviridae morphology, and **d)** Tolp, a *tectiviridae*. The scale bar for each grouping is shown below the image set.

### 3.4 HtPIP enables the discovery of the first *Levivirus* for *Microbacterium*

The twelfth novel phage discovered using HtPIP was *Microbacterium* phage Later isolated on *Microbacterium* sp.. This phage is a ssRNA virus in the *Leviviricetes* class with a 3,939nt genome. The closest relative based on nucleotide sequence is a *Riboviria sp*. isolate parrot82 MAG (OQ424403.1; 79.11%ID over 56% of the genome). To further characterize the phage genome we used VIPtree (33) to find related phages based on protein similarity, which revealed homology with common *Leviviruses* such as MS2, Qbeta, PP7, and AP205 (**Figure 5a**). While these phages share protein similarity, phage Later cannot be placed into phage taxonomy further than the family *Steitzviridae* based on coat protein analysis (46). *Microbacterium* phage Later encodes proteins typically found in *Leviviridae* genomes (47, 48), the maturation protein, coat protein, and replicase (**Figure 5b**). The discovery of *Microbacterium* phage Later demonstrates HtPIPs ability to discover RNA phages from environmental samples.

**Figure 5.**
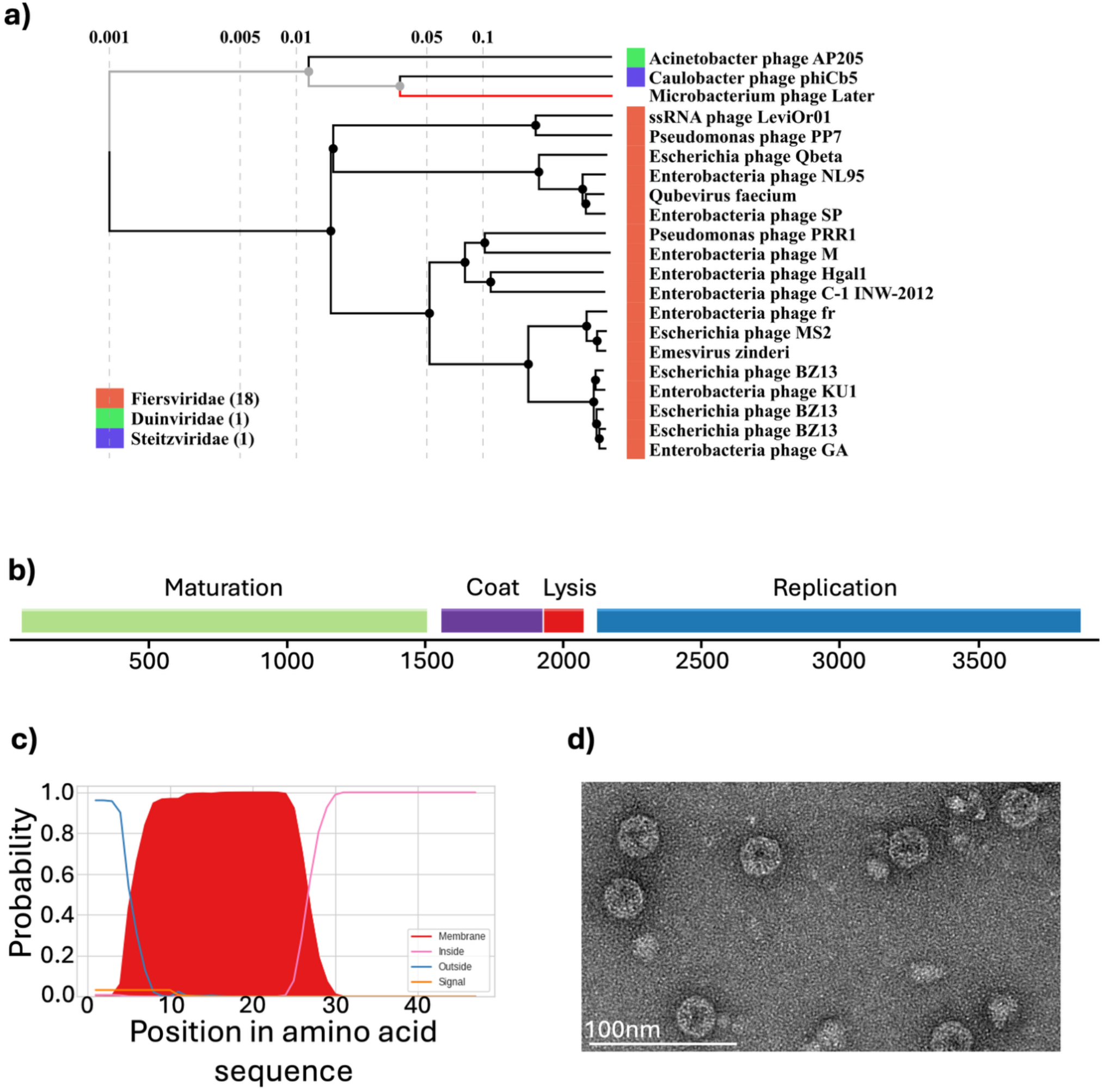
Isolation and Characterization of *Microbacterium* phage Later. **a)** VIP tree output for *Microbacterium* phage Later. Later appears to clade with the non-*Fiersviridae* in the VIP tree database. **b)** The annotated genome of Later. Protein functions are listed above the gene. **c)** deepTMHMM posterior probabilities for orf3 shows one transmembrane region. **d)** TEM of Later. The scale bar is shown in the lower left corner.

Since no *Leviviridae* have previously been described for *Actinobacteria* and standard annotation tools could not identify the lysis protein (which is not unexpected for *Leviviricetes* due to low sequence homology (47, 48)) we applied previously established criteria (48) to identify the lysis protein in *Microbacterium* phage Later. There was one predicted ORF that was annotated as a hypothetical protein (orf3). It was located between the coat protein and replication protein, a common location for *Leviviricetes* lysis proteins (**Figure 5b**). We analyzed the predicted ORF for transmembrane domains (TMD) and discovered a single TMD (**Figure 5c**), similar to other *Leviviridae* lysis proteins (48).

Lastly, to confirm *Microbacterium* phage Later was a *Leviviridae* we used TEM. We see small capsids 28nm x 25.4nm and no tail (**Figure 5d**). This morphology is the same as well-studied *Leviviricetes* that infect *Enterobacteria*, confirming *Microbacterium* phage Later is a *Leviviridae* and the first reported to infect *Actinobacteria* species.

### 3.5 Phage Diversity between HtPIP and Traditional Methods

While isolating novel phages showed that this method was compatible with a variety of environmental samples and bacteria, we were curious to see if the HtPIP captured different types of phages compared to traditional methods. We used *E. coli* MG1655 and a local wastewater influent sample for discovery since we observed total plate clearings in plaque assays from our local wastewater samples on *E. coli* for the first two rounds of phage hunting with HtPIP (**Supplemental Table 2**). We collected the virus fraction after initial enrichment from each method, extracted and sequenced the metaviromic DNA, and assembled into contigs >10kbp. Overall, 45 unique viral contigs were assembled between all sampling methods tested in this study.

We used MVP (38) to generate ORF predictions and the resultant protein sequences. Predicted proteins from the metaviromes we collected were clustered with VContact2 (39), which grouped the metaviromes into 10 unique viral clusters with at least two members among the 3 different sampling types, along with 18 singleton viral contigs (**Figure 6a**). Each cluster contained at least one member obtained with traditional methods (**Figure 6a**), which indicates that the traditional method captured phages that were similar variants to each other. The HtPiP Liquid and HtPiP Semisolid methods each yielded 6 singleton phage contigs (12 in total) compared to the 5 captured by traditional methods, suggesting they captured a distinct array of phages compared to traditional techniques (**Figure 6a**).

**Figure 6:**
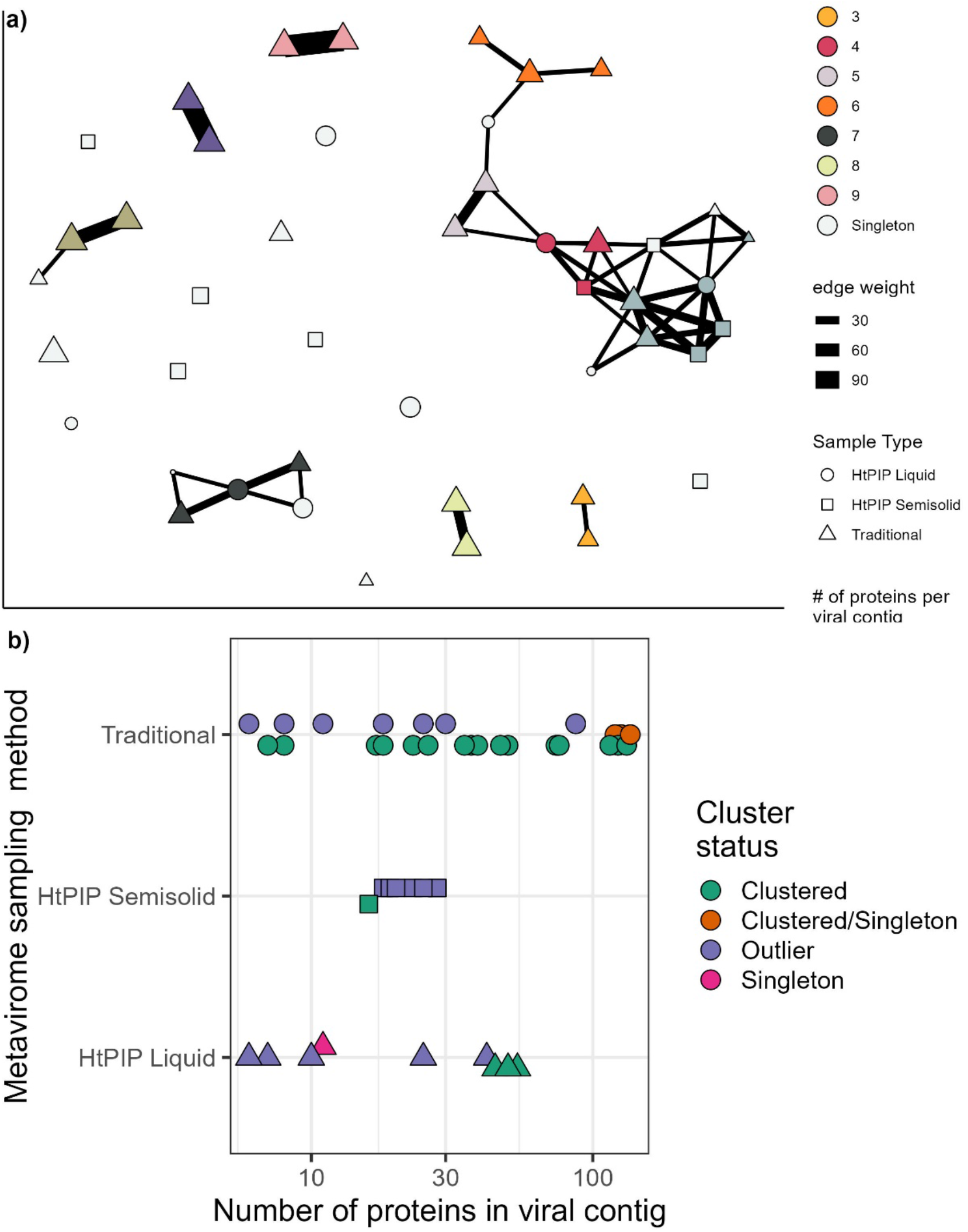
HtPIP loaded with liquid and semisolid media yields a higher proportion of novel viral contigs compared to traditional phage discovery. **a)** Intra-study viral contig clustering via VContact2. “Traditional” represents a filtered wastewater influent sample mixed with overnight *E. coli* diluted in LB and shaken for 3 days at 20ºC. The “HtPIP” samples originate from the HtPIP described in this paper. “Liquid” was when wells we filled with liquid LB, and “Solid” was when wells were filled with 0.5% LB Agar. Clusters were determined by VContact2. “Singleton” samples were phage contigs with no discernable neighbor within the metaviromes assembled in this study. **b)** Clustering phage contigs found in this study against NCBI RefSeq database via VContact2. The x-axis denotes the number of proteins identified per contig, which typically indicates the size of the viral genome. Shapes labelled “Clustered” represent an assembled viral contig with one or more close neighbors among the RefSeq database, suggesting a shared genus-level classification with a previously sequenced phage. “Clustered/Singleton” contigs shared some DNA segments that were clustered strongly and other segments that were not. “Outlier” contigs represent assembled viral contigs which clustered weakly with a known phage (based on edge weight), possibly representing a new genus. “Singleton” contigs did not cluster with any sequenced phage.

To determine the novelty of the *E. coli* phages recovered with each phage discovery technique, we next clustered the viral contigs against the known repository of phages found in NCBI RefSeq. For this analysis, viral contigs were clustered to other known viral contigs and classified as “Clustered”, “Outlier”, or “Singleton”. Clustered samples have significant homology to known phages in RefSeq but may be a new strain or species. Outlier samples have weak similarity to known phages and are possibly a new genus. Singleton contigs have no relatives at all and are the most unique category. Both HtPIP liquid and agar yielded a high proportion of outlier contigs to clustered contigs (85%, 63%, respectively) compared to traditional methods (27%) (**Figure 6b**), suggesting that HtPIP captures a higher proportion of previously undiscovered phages compared to traditional methods.

## Discussion

In this study, we have developed a novel high-throughput approach to discover phages simultaneously for multiple bacterial hosts and tested it with a variety of environmental samples. We isolated twelve new phages for nine different bacterial hosts. Many of these phages are genomically and morphologically novel compared to previously described phages and cannot be taxonomically classified beyond the family level. Within this set of novel phages, we found both DNA and RNA phages, multiple morphotypes, and many new predicted gene products. Additionally, we compared metaviromes obtained using our method versus traditional methods and found that HtPIP captures a distinct set of phages compared to traditional methods (**Figure 6a**), and a higher proportion of previously undiscovered/outlier phages than traditional methods (**Figure 6b**). Importantly, the HtPIP technique saves time and requires simpler equipment for phage discovery, specifically during the initial filtering step of environmental samples; this step can involve up to 3 hours of centrifuging, incubation, and filtering (22) and is not performed in our method.

The HtPIP is especially amenable to search for phages across large-scale host collections, as the user can aliquot overnight cultures of a culture collection into multiple plates during the initial setup phase and incubate these replicate plates on top of several distinct environmental samples. For this study, we set up triplicate plates and placed one replicate plate on top of one of three distinct environmental samples. The 96 well format is also compatible with automated lab equipment such as liquid handlers and colony pickers and downstream approaches to high-throughput phage isolation (19). Though not explored in this study, this system could also be used in a time-series manner in environments where the phage ecology of the sample changes with time, as in soil samples (49, 50). Phage discovery from soil samples or other solid samples will especially benefit from using the HtPIP since filtering these sample types usually entails multiple centrifugation steps and prefiltration (22, 23). Further it is possible to adapt HtPIP to work *in situ* without risk of contaminating nearby soils or water since sealed plates are used.

While the HtPIP allowed for discovery of many novel phages and provides a platform to speed phage discovery, there are limitations. One limitation is that some bacterial hosts seem to be more difficult to find phages for. We searched for *Burkholderia* phages in seven different samples with no success. While not published in this article, we searched a similar number of samples for the same *Burkholderia* host with traditional methods and found no phages. This seems consistent with the few phages described in the literature for *Burkholderia*, so different techniques are likely needed to isolate phages for *Burkholderia* (51). The second limitation is downstream screening, in our current work we still performed double-agar overlays to identify phage plaques after enrichment with the HtPIP. This step could be accelerated in the future by using automated lab equipment and workflows. Finally, our current HtPIP setup has a single reservoir, so only one environmental sample can be screened across up to 96 bacterial strains. While this significantly enhances the phage discovery process, many workflows will not require this many bacterial strains to screen against. Custom plates can be developed to allow for multiple reservoirs to potentially screen multiple environmental samples against fewer bacterial hosts, or these plates may be cut in half with a precision tool.

While phage discovery has accelerated in recent years (52, 53) due to a renewed interest in phages for medical, health, and biotechnology applications (54), many bacteria still have few or no phage isolates (52, 55). The HtPIP enables researchers to place up to 96 different bacteria into the plate with each sample which increases the chance that a phage for a non-model bacteria may be found in an environmental sample. Discovery of phage isolates for non-model bacteria, especially those gaining interest as biomanufacturing chassis, provides valuable toolkits for these bacteria. These phages can be used to deliver gene cassettes to previously difficult to transform bacteria or bacteria in microbial communities or the phage gene products can be used to further enhance biotechnology applications (10, 11).

One interesting discovery from our new phages was the extremely long tails found on the *Rhodococcus* phages. Maizie, SweetSinh, Prosser1, and Perlinasted all have tails >400nm which are among some of the longest tails found on isolated phages that can be repeatedly infected. There is one phage described for *R. equi*, ReqiDocB7 which also has a 489nm tail (56). Very long flexible tails may be necessary to go through the thick cell wall of *Rhodococcus* (57).

The HtPIP enabled discovery of new *Tectiviridae* (**Figure 4**), which can be difficult to isolate with traditional methods. *Tectiviridae* are known to infect *Pseudomonas* species which carry plasmids for pili formation (58, 59). We confirm here our new phage Tolp can infect *P. putida* mt-2, which harbors the pWW0 plasmid that contains genes for conjugative transfer and toluene and xylene degradation, (60, 61), but not *P. putida KT2440*, which lacks this plasmid (**Supplementary Table 2**).

The most taxonomically novel phage found using the HtPIP was the RNA phage Later, which we confirmed through genomic sequencing and TEM to be a *Leviviricete* infecting a soil *Microbacterium* isolate (**Figure 5**). This demonstrates the ability of HtPIP to capture both DNA and RNA viruses, and is the first RNA phage isolated outside of proteobacteria (62). The only other RNA phage described outside of proteobacteria is *Streptomyces* phage ϕ0, a *Cystovirdae*, uncovered in a pure culture transcriptome experiment from *S. avermitilis* (46, 62). Although *Leviviridae* isolates have been described for many gram-negative hosts including, *Enterobacteria, Pseudomonas, Acinetobacter*, and *Caulobacter*, to date there has been no *Leviviridae* isolated for *Actinobacteria* (46, 63-66) or any gram-positive host to our knowledge. Many outstanding questions remain to be explored for Later, especially regarding phage-host interactions required for entry. All *Leviviridae* described use Gram-negative bacterial pili to begin the infection process (67). While our *Microbacterium* isolate has an annotated Type IV pilus assembly operon, it is unknown if the mechanism of phage entry is different for Gram-positive bacteria. It’s also possible quorum signaling agents or antibiotics produced by the native bacterial community growing in the reservoir beneath the semipermeable plate may promote the expression of surface structures similar to pili (68) that are critical for *Leviviridae* attachment and subsequent infection. These and possibly other unknown chemicals exchanged between the growing native bacterial community and the monoculture hosts grown on the other side of the membrane warrants further research to better understand how the microbiome community affects phage infection dynamics.

## Conclusion

Here we present a novel phage discovery platform (HtPIP), which grows bacterial hosts in close proximity to environmental samples such as soil and sewage influent, separated by a 0.2-micron membrane. We optimized a procedure to capture the most phages using model bacterial hosts and tested the HtPIP across a variety of environmental samples to discover phages for multiple hosts at once. Using this method, we were able to capture DNA and RNA phages infecting a diverse range of hosts, including both gram negative and positive bacteria. We also compare the diversity and novelty of phages from the metaviromes of enrichment cultures and found that a higher proportion of unique phage contigs were assembled from HtPIP compared to traditional phage discovery techniques. We envision researchers using HtPIP to efficiently discover diverse and novel phages for up to 96 bacteria in a single sample.

## Author Contributions

BD: Conceptualization, Investigation, Methodology, Visualization, Writing – original draft; Writing – review & editing; TH: Investigation, Writing – review & editing; MP: Investigation, Writing – review & editing; JSS: Funding acquisition, Writing – review & editing; CMM: Conceptualization, Investigation, Data curation, Supervision, Funding acquisition, Visualization, Writing – original draft; Writing – review & editing;

## Acknowledgments

We thank John Becky and Rodolphe Barrangou (North Carolina State University) for sequencing the Soil63 phage genome at SeqCoast Genomics and kindly sharing *Variovorax sp*. OAS795 and *Variovorax sp*. SCN. We thank Ryan McClure and Kirsten Hofmockel (Pacific Northwest National Laboratory) for sharing the MSC-2 isolates, including *Rhodococcus sp*. MSC1_016, *Sphingopyxis fribergensis* MSC1_008, and *Microbacterium sp*.. We thank Brady Cress (Integrative Genomics Institute) for sharing *B. subtilis* NCIB3610. We thank Brady Cress, Michael Cohen, and Irina Khilyas for providing *Rhodococcus qingshengii* S10.

All TEM imagining was done by Tagide deCarvalho, Keith R. Porter Imaging Facility, UMBC.

We thank Taylor Moehling for internal review of this manuscript.

The following reagent was obtained through BEI Resources, NIAID, NIH: *Burkholderia cenocepacia*, strain K56-2 (Valvano), NR-20535.

## Data Availability Statement

The metaviromics datasets generated in this study can be found in the sequence read archive (SRA) under BioProject PRJNA1415247, Biosamples SAMN54927485-SAMN54927489 and SAMN54985761. All phage genomes were deposited into NCBI GenBank, accession numbers are listed in **Supplemental Table 1**.

## References

1. Abe K, Kawano Y, Iwamoto K, Arai K, Maruyama Y, Eichenberger P, et al. Developmentally-regulated excision of the SPβ prophage reconstitutes a gene required for spore envelope maturation in Bacillus subtilis. PLoS Genet. 2014;10(10):e1004636.

2. Graham EB, Camargo AP, Wu R, Neches RY, Nolan M, Paez-Espino D, et al. A global atlas of soil viruses reveals unexplored biodiversity and potential biogeochemical impacts. Nat Microbiol. 2024;9(7):1873–83.

3. Eppley JM, Biller SJ, Luo E, Burger A, DeLong EF. Marine viral particles reveal an expansive repertoire of phage-parasitizing mobile elements. Proc Natl Acad Sci U S A. 2022;119(43):e2212722119.

4. Shkoporov AN, Turkington CJ, Hill C. Mutualistic interplay between bacteriophages and bacteria in the human gut. Nat Rev Microbiol. 2022;20(12):737–49.

5. Mageeney CM, Mohammed HT, Dies M, Anbari S, Cudkevich N, Chen Y, et al. Mycobacterium Phage Butters-Encoded Proteins Contribute to Host Defense against Viral Attack. mSystems. 2020;5(5).

6. Dedrick RM, Jacobs-Sera D, Bustamante CA, Garlena RA, Mavrich TN, Pope WH, et al. Prophage-mediated defence against viral attack and viral counter-defence. Nat Microbiol. 2017;2:16251.

7. Bondy-Denomy J, Qian J, Westra ER, Buckling A, Guttman DS, Davidson AR, et al. Prophages mediate defense against phage infection through diverse mechanisms. Isme j. 2016;10(12):2854–66.

8. Wu R, Smith CA, Buchko GW, Blaby IK, Paez-Espino D, Kyrpides NC, et al. Structural characterization of a soil viral auxiliary metabolic gene product - a functional chitosanase. Nat Commun. 2022;13(1):5485.

9. Karimi M, Mirshekari H, Moosavi Basri SM, Bahrami S, Moghoofei M, Hamblin MR. Bacteriophages and phage-inspired nanocarriers for targeted delivery of therapeutic cargos. Adv Drug Deliv Rev. 2016;106(Pt A):45–62.

10. Lammens EM, Nikel PI, Lavigne R. Exploring the synthetic biology potential of bacteriophages for engineering non-model bacteria. Nat Commun. 2020;11(1):5294.

11. Lemire S, Yehl KM, Lu TK. Phage-Based Applications in Synthetic Biology. Annu Rev Virol. 2018;5(1):453–76.

12. Nakayinga R, Makumi A, Tumuhaise V, Tinzaara W. Xanthomonas bacteriophages: a review of their biology and biocontrol applications in agriculture. BMC Microbiol. 2021;21(1):291.

13. Huss P, Raman S. Engineered bacteriophages as programmable biocontrol agents. Curr Opin Biotechnol. 2020;61:116–21.

14. Figueiredo CM, Malvezzi Karwowski MS, da Silva Ramos RCP, de Oliveira NS, Peña LC, Carneiro E, et al. Bacteriophages as tools for biofilm biocontrol in different fields. Biofouling. 2021;37(6):689–709.

15. Hatfull GF, Dedrick RM, Schooley RT. Phage Therapy for Antibiotic-Resistant Bacterial Infections. Annu Rev Med. 2022;73:197–211.

16. Gordillo Altamirano FL, Barr JJ. Phage Therapy in the Postantibiotic Era. Clin Microbiol Rev. 2019;32(2).

17. Weber-Dąbrowska B, Jończyk-Matysiak E, Żaczek M, Łobocka M, Łusiak-Szelachowska M, Górski A. Bacteriophage Procurement for Therapeutic Purposes. Front Microbiol. 2016;7:1177.

18. Hyman P. Phages for Phage Therapy: Isolation, Characterization, and Host Range Breadth. Pharmaceuticals (Basel). 2019;12(1).

19. Olsen NS, Hendriksen NB, Hansen LH, Kot W. A New High-Throughput Screening Method for Phages: Enabling Crude Isolation and Fast Identification of Diverse Phages with Therapeutic Potential. Phage (New Rochelle). 2020;1(3):137–48.

20. Nichols D, Cahoon N, Trakhtenberg EM, Pham L, Mehta A, Belanger A, et al. Use of ichip for high-throughput in situ cultivation of “uncultivable” microbial species. Appl Environ Microbiol. 2010;76(8):2445–50.

21. Wu R, Davison MR, Nelson WC, Smith ML, Lipton MS, Jansson JK, et al. Hi-C metagenome sequencing reveals soil phage-host interactions. Nat Commun. 2023;14(1):7666.

22. Göller PC, Haro-Moreno JM, Rodriguez-Valera F, Loessner MJ, Gómez-Sanz E. Uncovering a hidden diversity: optimized protocols for the extraction of dsDNA bacteriophages from soil. Microbiome. 2020;8(1):17.

23. Trubl G, Roux S, Solonenko N, Li YF, Bolduc B, Rodríguez-Ramos J, et al. Towards optimized viral metagenomes for double-stranded and single-stranded DNA viruses from challenging soils. PeerJ. 2019;7:e7265.

24. Brauer A, Rosendahl S, Kängsep A, Lewańczyk AC, Rikberg R, Hõrak R, et al. Isolation and characterization of a phage collection against Pseudomonas putida. Environ Microbiol. 2024;26(6):e16671.

25. Bankevich A, Nurk S, Antipov D, Gurevich AA, Dvorkin M, Kulikov AS, et al. SPAdes: a new genome assembly algorithm and its applications to single-cell sequencing. J Comput Biol. 2012;19(5):455–77.

26. Seemann T. Prokka: rapid prokaryotic genome annotation. Bioinformatics. 2014;30(14):2068–9.

27. Chan PP, Lin BY, Mak AJ, Lowe TM. tRNAscan-SE 2.0: improved detection and functional classification of transfer RNA genes. Nucleic Acids Res. 2021;49(16):9077–96.

28. Millard A, Denise R, Lestido M, Thomas MT, Webster D, Turner D, et al. taxMyPhage: Automated Taxonomy of dsDNA Phage Genomes at the Genus and Species Level. Phage (New Rochelle). 2025;6(1):5–11.

29. Bolduc B, Jang HB, Doulcier G, You ZQ, Roux S, Sullivan MB. vConTACT: an iVirus tool to classify double-stranded DNA viruses that infect Archaea and Bacteria. PeerJ. 2017;5:e3243.

30. Arkin AP, Cottingham RW, Henry CS, Harris NL, Stevens RL, Maslov S, et al. KBase: the United States department of energy systems biology knowledgebase. Nature biotechnology. 2018;36(7):566–9.

31. Grant JR, Enns E, Marinier E, Mandal A, Herman EK, Chen CY, et al. Proksee: in-depth characterization and visualization of bacterial genomes. Nucleic Acids Res. 2023;51(W1):W484–w92.

32. Nayfach S, Camargo AP, Schulz F, Eloe-Fadrosh E, Roux S, Kyrpides NC. CheckV assesses the quality and completeness of metagenome-assembled viral genomes. Nat Biotechnol. 2021;39(5):578–85.

33. Nishimura Y, Yoshida T, Kuronishi M, Uehara H, Ogata H, Goto S. ViPTree: the viral proteomic tree server. Bioinformatics. 2017;33(15):2379–80.

34. Moraru C, Varsani A, Kropinski AM. VIRIDIC-A Novel Tool to Calculate the Intergenomic Similarities of Prokaryote-Infecting Viruses. Viruses. 2020;12(11).

35. Li D, Luo R, Liu CM, Leung CM, Ting HF, Sadakane K, et al. MEGAHIT v1.0: A fast and scalable metagenome assembler driven by advanced methodologies and community practices. Methods. 2016;102:3–11.

36. Li D, Liu CM, Luo R, Sadakane K, Lam TW. MEGAHIT: an ultra-fast single-node solution for large and complex metagenomics assembly via succinct de Bruijn graph. Bioinformatics. 2015;31(10):1674–6.

37. Hallgren J, Tsirigos KD, Pedersen MD, Almagro Armenteros JJ, Marcatili P, Nielsen H, et al. DeepTMHMM predicts alpha and beta transmembrane proteins using deep neural networks. bioRxiv. 2022:2022.04.08.487609.

38. Coclet C, Camargo AP, Roux S. MVP: a modular viromics pipeline to identify, filter, cluster, annotate, and bin viruses from metagenomes. mSystems. 2024;9(10):e0088824.

39. Bin Jang H, Bolduc B, Zablocki O, Kuhn JH, Roux S, Adriaenssens EM, et al. Taxonomic assignment of uncultivated prokaryotic virus genomes is enabled by gene-sharing networks. Nat Biotechnol. 2019;37(6):632–9.

40. Jaryenneh J, Schoeniger JS, Mageeney CM. Genome sequence and characterization of a novel Pseudomonas putida phage, MiCath. Sci Rep. 2023;13(1):21834.

41. Bucher MJ, Czyż DM. Phage against the Machine: The SIE-ence of Superinfection Exclusion. Viruses. 2024;16(9).

42. Jaryenneh J, Krishna R, Schoeniger JS, Mageeney CM. Complete genome sequence of Rhodococcus qingshengii phage Perlina. Microbiol Resour Announc. 2024;13(11):e0086924.

43. McClure R, Farris Y, Danczak R, Nelson W, Song HS, Kessell A, et al. Interaction Networks Are Driven by Community-Responsive Phenotypes in a Chitin-Degrading Consortium of Soil Microbes. mSystems. 2022;7(5):e0037222.

44. Coutinho FH, Silveira CB, Sebastián M, Sánchez P, Duarte CM, Vaqué D, et al. Water mass age structures the auxiliary metabolic gene content of free-living and particle-attached deep ocean viral communities. Microbiome. 2023;11(1):118.

45. Hutinet G, Kot W, Cui L, Hillebrand R, Balamkundu S, Gnanakalai S, et al. 7-Deazaguanine modifications protect phage DNA from host restriction systems. Nat Commun. 2019;10(1):5442.

46. Callanan J, Stockdale SR, Adriaenssens EM, Kuhn JH, Rumnieks J, Pallen MJ, et al. Leviviricetes: expanding and restructuring the taxonomy of bacteria-infecting single-stranded RNA viruses. Microb Genom. 2021;7(11).

47. Ruokoranta TM, Grahn AM, Ravantti JJ, Poranen MM, Bamford DH. Complete genome sequence of the broad host range single-stranded RNA phage PRR1 places it in the Levivirus genus with characteristics shared with Alloleviviruses. J Virol. 2006;80(18):9326–30.

48. Chamakura KR, Tran JS, O’Leary C, Lisciandro HG, Antillon SF, Garza KD, et al. Rapid de novo evolution of lysis genes in single-stranded RNA phages. Nat Commun. 2020;11(1):6009.

49. Wu R, Davison MR, Gao Y, Nicora CD, McDermott JE, Burnum-Johnson KE, et al. Moisture modulates soil reservoirs of active DNA and RNA viruses. Commun Biol. 2021;4(1):992.

50. Van Goethem MW, Swenson TL, Trubl G, Roux S, Northen TR. Characteristics of Wetting-Induced Bacteriophage Blooms in Biological Soil Crust. mBio. 2019;10(6).

51. Lauman P, Dennis JJ. Advances in Phage Therapy: Targeting the Burkholderia cepacia Complex. Viruses. 2021;13(7).

52. Cook R, Brown N, Redgwell T, Rihtman B, Barnes M, Clokie M, et al. INfrastructure for a PHAge REference Database: Identification of Large-Scale Biases in the Current Collection of Cultured Phage Genomes. Phage (New Rochelle). 2021;2(4):214–23.

53. Shen J, Zhang J, Mo L, Li Y, Li Y, Li C, et al. Large-scale phage cultivation for commensal human gut bacteria. Cell Host Microbe. 2023;31(4):665-77.e7.

54. Santini JM. The new age of the phage. Essays Biochem. 2024;68(5):579–81.

55. Roux S, Mutalik VK. Tapping the treasure trove of atypical phages. Curr Opin Microbiol. 2024;82:102555.

56. Summer EJ, Liu M, Gill JJ, Grant M, Chan-Cortes TN, Ferguson L, et al. Genomic and functional analyses of Rhodococcus equi phages ReqiPepy6, ReqiPoco6, ReqiPine5, and ReqiDocB7. Appl Environ Microbiol. 2011;77(2):669–83.

57. Piselli C, Benier L, Koy C, Glocker MO, Benz R. Cell wall channels of Rhodococcus species: identification and characterization of the cell wall channels of Rhodococcus corynebacteroides and Rhodococcus ruber. Eur Biophys J. 2022;51(4-5):309–23.

58. Grahn AM, Butcher SJ, Bamford JK, Bamford DH. PRD1: dissecting the genome, structure and entry. The bacteriophages. 2006;2(2677):161.

59. Saren AM, Ravantti JJ, Benson SD, Burnett RM, Paulin L, Bamford DH, et al. A snapshot of viral evolution from genome analysis of the tectiviridae family. J Mol Biol. 2005;350(3):427–40.

60. Burlage RS, Hooper SW, Sayler GS. The TOL (pWW0) catabolic plasmid. Appl Environ Microbiol. 1989;55(6):1323–8.

61. Greated A, Lambertsen L, Williams PA, Thomas CM. Complete sequence of the IncP-9 TOL plasmid pWW0 from Pseudomonas putida. Environ Microbiol. 2002;4(12):856–71.

62. Krishnamurthy SR, Janowski AB, Zhao G, Barouch D, Wang D. Hyperexpansion of RNA Bacteriophage Diversity. PLoS Biol. 2016;14(3):e1002409.

63. Kazaks A, Voronkova T, Rumnieks J, Dishlers A, Tars K. Genome structure of caulobacter phage phiCb5. J Virol. 2011;85(9):4628–31.

64. Fiers W, Contreras R, Duerinck F, Haegeman G, Iserentant D, Merregaert J, et al. Complete nucleotide sequence of bacteriophage MS2 RNA: primary and secondary structure of the replicase gene. Nature. 1976;260(5551):500–7.

65. Klovins J, Overbeek GP, van den Worm SHE, Ackermann HW, van Duin J. Nucleotide sequence of a ssRNA phage from Acinetobacter: kinship to coliphages. J Gen Virol. 2002;83(Pt 6):1523–33.

66. Kim ES, Bae HW, Cho YH. A Pilin Region Affecting Host Range of the Pseudomonas aeruginosa RNA Phage, PP7. Front Microbiol. 2018;9:247.

67. Hör J. Advancing RNA phage biology through meta-omics. Nucleic Acids Res. 2025;53(8).

68. Lu Y, Zeng J, Wang L, Lan K E S, Wang L, et al. Antibiotics Promote Escherichia coli-Pseudomonas aeruginosa Conjugation through Inhibiting Quorum Sensing. Antimicrob Agents Chemother. 2017;61(12).

