## Supplementary File for "HtPIP: High-Throughput Phage Isolation Platform increases diversity and reduces isolation time using multiple bacteria"

### Supplementary Materials

**Supplemental Table 1. Collection information, genotypic, and phenotypic information about all isolated phages.**

| | **Wastewater Influent 1** | **Compost - backyard** | **Wastewater influent 2** | **Commercial Topsoil** | **Tomato rhizosphere** | **Wastewater influent 3** | **Compost - community center** | **Agricultural soil** |
|---|---|---|---|---|---|---|---|---|
| *E. coli* MG1655 | | | | | | | | |
| *P. putida* S12 | | | | | | | | |
| *P. putida* mt-2 | | | | | | | | |
| *B. cenocepacia K*56-2 | | | | | | | | |
| *Microbacterium* sp. MSC1_018 | | | | | | | | |
| *R. rhodochrous* 372 | | | | | | | | |
| *R. qingshengii S10* | | | | | | | | |
| *Variovorax sp. SCN* | | | | | | | | |
| *Variovorax* sp. OAS795 | | | | | | | | |
| *S. fribergensis* MSC1_008 | | | | | | | | |
| *Rhodococcus sp.* MSC1_016 | | | | | | | | |

**Supplementary Table 2. Environmental Sample and Bacterial Host Matrix.** Light blue boxes denote plaques isolated from enrichments; dark blue denotes web of phages were isolated. White denotes no phages were found. Grey boxes denote not tested. The total combinations of strains and samples tested was 54.
