## Supplemental Table 1 for "HtPIP: High-Throughput Phage Isolation Platform increases diversity and reduces isolation time using multiple bacteria"

| Phage Name | Bacterial Host | Environmental Sample | Sequencing Platform (Vendor) | DNA extraction method | Genbank Accession Number | Genome Length (bp/nt) | %GC content | Coverage | Protein-coding genes (tRNAs) | Proteins with predicted functions | Taxonomy | Morphotype | Virion head diameter (nm) | Virion tail length (nm) |
|---|---|---|---|---|---|---|---|---|---|---|---|---|---|---|
| EW1 | *E. coli* MG1655 | Wastewater Influent 1 | MiSeq (SNL) | Norgen | PV920661 | 49619 | 45.6 | 2133 | 81 | 40 | Genus:Tunavirus | Siphovirdae | 45 | 150 |
| Eliverwaste | *E. coli* MG1655 | Wastewater Influent 1 | MiSeq (SNL) | Norgen | PV933152 | 88023 | 39.1 | 1139 | 136 (22) | 51 | Genus: Felixounavirus | Myoviridae | 60 | 100 |
| PLiverWaste | *P. putida* mt-2/ KT2440 | Wastewater Influent 2 | MiSeq (SNL) | Norgen | PV933153 | 53523 | 54.9 | 360 | 75 | 31 | Class: Caudoviricetes | Siphovirdae | 60 | 143 |
| Tolp | *P. putida* mt-2 | Wastewater Influent 2 | MiSeq (SNL) | Norgen | PV920658 | 14926 | 48.3 | 330 | 30 | 26 | Species: Alphatectivirus PRD1 | Tectivirdae | 60 | 40 |
| Sweet Sinh | *R. rhodochrous* 372 | Commercial Topsoil | MiSeq (SNL) | Norgen | PV933154 | 78423 | 65.4 | 38 | 101 | 35 | Class: Caudoviricetes | Siphovirdae | 60 | 400 |
| Perlinasted | *R. qingshengii* S10 | Compost - community center | MiSeq (SNL) | Norgen | PV920656 | 75747 | 54.1 | 23 | 101 (3) | 40 | Class: Caudoviricetes | Siphovirdae | 60 | 440 |
| Maizie | *R. rhodochrous* 372 | Commercial Topsoil | MiSeq (SNL) | Norgen | PV933155 | 78509 | 65.5 | 23 | 102 | 36 | Class: Caudoviricetes | Siphovirdae | 60 | 400 |
| Prosser1 | *Rhodococcus sp.* MSC1_016 | Agricultural soil | Illumina (SeqCoast) | Bead-beating lysis (SeqCoast Genomics) | PV920660 | 80548 | 56.9 | 3425 | 107 (1) | 35 | Class: Caudoviricetes | Siphovirdae | 60 | 460 |
| Gard | *Variovorax sp.* SCN | Compost - community center | MiSeq (SNL) | Norgen | PV920657 | 43634 | 62.3 | 131 | 58 | 29 | Family: Autographiviridae | Myoviridae | 90 | 200 |
| Soil63 | *Variovorax sp.* OAS795 | Agricultural soil | Illumina (SeqCoast) | Bead-beating lysis (SeqCoast Genomics) | PV933156 | 103377 | 61.4 | 1723.4 | 141 | 32 | Class: Caudoviricetes | Siphovirdae | 55 | 230.7 |
| Prosser2 | *S. fribergensis* MSC1_008 | Agricultural soil | Illumina (SeqCoast) | Bead-beating lysis (SeqCoast Genomics) | PV920659 | 62621 | 59.9 | 2277 | 82 | 40 | Class: Caudoviricetes | Siphovirdae | 54.8nmx28.6nm | 166 |
| Later | *Microbacterium sp.* MSC18 | Compost - community center | NextSeq2000 (SNL) | Monarch Mag Viral RNA/DNA extraction kit | PX757644 | 3939 | 49.38 | 58706.839 | 4 | 4 | Family:Steitzviridae | Levivirdae | 28nm x 25.4nm | N/A |
